# Crosstalk between guanosine nucleotides regulates cellular heterogeneity in protein synthesis during nutrient limitation

**DOI:** 10.1101/2021.11.22.469502

**Authors:** Simon Diez, Molly Hydorn, Abigail Whalen, Jonathan Dworkin

## Abstract

Phenotypic heterogeneity of microbial populations can facilitate survival in dynamic environments by generating sub-populations of cells that may have differential fitness in a future environment. *Bacillus subtilis* cultures experiencing nutrient limitation contain distinct sub-populations of cells exhibiting either comparatively high or low protein synthesis activity. This heterogeneity requires the production of phosphorylated guanosine nucleotides (pp)ppGpp by three synthases: SasA, SasB, and RelA. Here we show that these enzymes differentially affect this bimodality: RelA and SasB are necessary to generate the sub-population of cells exhibiting low protein synthesis whereas SasA is necessary to generate cells exhibiting comparatively higher protein synthesis. The RelA product (pppGpp) allosterically activates SasB and we find, in contrast, that the SasA product (pGpp) competitively inhibits this activation. Finally, we provide *in vivo* evidence that this antagonistic interaction mediates the observed heterogeneity in protein synthesis. This work therefore identifies the mechanism underlying phenotypic heterogeneity in the central physiological process of protein synthesis.

**Author Summary:** Upon encountering conditions that are unfavorable to growth, such as nutrient limitation, bacteria enter into a quiescent phenotype that is mediated by group of guanosine nucleotides collectively known as (pp)pGpp. These nucleotides direct the down-regulation of energy intensive processes and are essential for a striking heterogeneity in protein synthesis observed during exit from rapid growth. Here, we show that a network of (pp)pGpp synthases is responsible for this heterogeneity and describe a mechanism that allows for the integration of multiple signals into the decision to down regulate the most energy intensive process in a cell.

## Introduction

Nutrient availability is a major environmental cue for bacteria. For example, amino acid starvation results in induction of the stringent response, a conserved mechanism dependent on the synthesis of the nucleotides guanosine penta- and tetra-phosphate ((p)ppGpp). These nucleotides mediate a broad shut down of energy intensive reactions which are required during rapid growth (1, 2). (p)ppGpp directly binds and inhibits key proteins that catalyze processes including transcription (RNA polymerase (3, 4)), translation (GTPase IF2 (5)), GTP biosynthesis (HprT and GmK (6)), DNA replication (DNA primase (7)), and ribosome assembly (ObgE and RsgA (8)).

Gram-positive bacteria typically encode a single, bi-functional RSH enzyme capable of both (p)ppGpp synthesis and hydrolysis as well as two additional small alarmone synthases (SAS) that lack hydrolytic activity. Unlike RSH proteins, which are activated by the binding of deacylated tRNAs to the A-site of the ribosome, SAS enzymes are believed to be transcriptionally regulated (9) and some are also under allosteric control (10). RelA/SpoT and the SAS synthases preferentially produce different molecules in different species. For example, in response to amino acid starvation, *E. coli* RelA produces approximately equal amounts of the tetra-phosphorylated (ppGpp) and the penta-phosphorylated (pppGpp) guanosines, whereas *B. subtilis* RelA primarily generates pppGpp using GTP and ATP as substrates (11). *B. subtilis* SasB preferentially utilizes GDP and ATP to generate the tetra-phosphorylated guanosine (ppGpp) (12) and SasA, the other SAS enzyme in *B. subtilis*, primarily generates a 5’ monophosphate 3’ di-phosphate guanosine (pGpp) using GMP and ATP as substrates *in vivo* (12). Together, these three closely related nucleotides are referred to as (pp)pGpp.

Recently, our laboratory demonstrated that accumulation of (pp)pGpp attenuates protein synthesis when populations of *B. subtilis* cease growing exponentially (5). This attenuation is bimodal and results in a heterogeneity in the protein synthesis activity of individual cells that exhibit either comparatively high or low protein synthesis activity (5). Here we find that a network of interacting (pp)pGpp synthases including a RSH protein (RelA) and two SAS proteins (SasA, SasB) underlies this heterogeneity since the absence of any of these synthases results in the loss of bimodality. The products of SasA and RelA, pGpp and pppGpp respectively, together antagonistically regulate activation of the third synthase (SasB), that is itself responsible for the synthesis of ppGpp, which inhibits Initiation Factor 2 and thereby attenuates protein synthesis (5).

## Results

### The SasA and SasB (p)ppGpp synthases contribute to heterogeneity

Cellular heterogeneity in protein synthesis as *B. subtilis* cultures exit rapid growth is dependent on the presence of the phosphorylated guanosine nucleotides (pp)pGpp (5). We investigated the origins of this heterogeneity by assessing single cell protein synthesis using *O*-propargyl-puromycin (OPP) incorporation in strains carrying deletion mutations in either of the two *B. subtilis* (pp)pGpp synthases (SasA and SasB) whose expression increases during exit from rapid growth (12). To quantify these effects we applied a cutoff that specifies the population of cells with low rates of protein synthesis. We set the threshold of this cutoff (850 relative fluorescence units (RFU)) as the magnitude of OPP labeling of wildtype *B. subtilis* culture in late transition phase (Fig. S1A) that captures 95% of the entire population. We used this threshold to define the fraction of the population with low rates of protein synthesis (“OFF”) (Fig. S1B). By convention, we define the remainder of the population as “ON.”

A strain lacking SasB (*ΔsasB*) contained fewer “OFF” cells as compared to the wildtype strain (Fig. 1A, B; S2). This result is consistent with our previous observation that the SasB product ppGpp inhibits the function of IF2 and thereby downregulates protein synthesis (5). In contrast, a strain lacking SasA (*ΔsasA*) does not contain the substantial fraction of “ON” cells seen in the wildtype parent strain (Fig. 1A, C; S2) and most cells in the population are “OFF”. This observation suggests that the SasA product pGpp does not directly inhibit translation, as does ppGpp, but rather acts indirectly.

**Fig 1.**
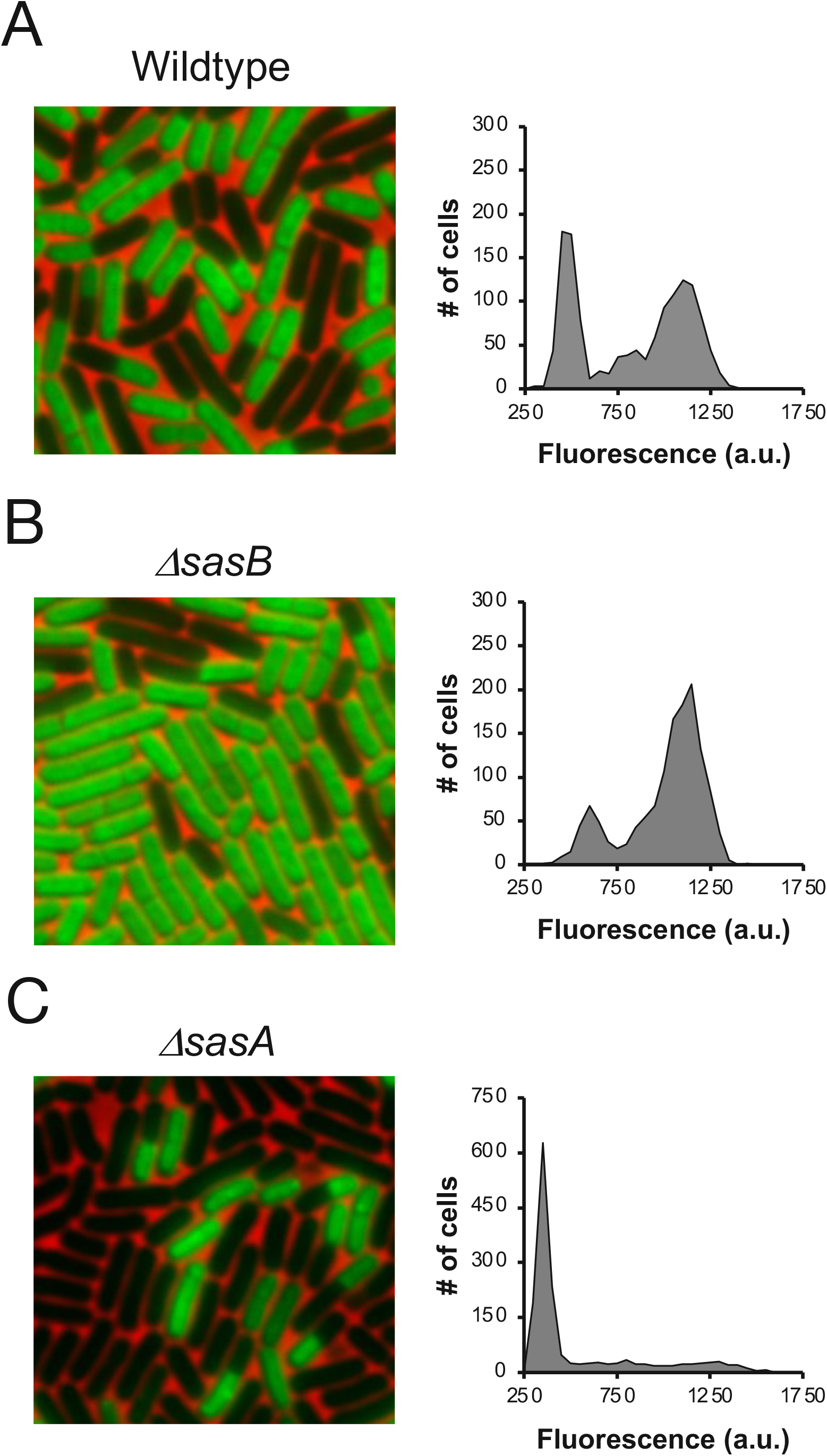
*sasB* and *sasA* have opposing effects on bimodality. **(A, B, C)** Representative pictures and population distributions of OPP labeled **(A)** wildtype (JDB1772), **(B)** Δ*sasB* (JDB4310) and **(C)** Δ*sasA* (JDB4311) during late transition phase.

### *sasA* but not *sasB* expression is correlated with levels of protein synthesis

*sasA* and *sasB* are regulated transcriptionally and expressed post-exponentially (12, 13) when the heterogeneity is observed (Fig 1A). We therefore asked if expression of either *sasA* or *sasB* is correlated with protein synthesis using transcriptional fusions of the *sasA* or the *sasB* promoters to YFP (*P*_*sasA*_*-yfp* or *P*_*sasB*_*-yfp)*. Consistent with prior observations (12), expression of both *sasA* and *sasB* reporters increased during the exit from exponential growth (Fig 2A, B). We examined the relationship between promoter activity and protein synthesis by measuring both YFP expression and OPP incorporation in single cells. Cells with higher *sasA* expression (*P*_*sasA*_*-yfp)* are more likely to have higher levels of protein synthesis than cells with lower *sasA* expression (Fig 2D). If the population is divided into quartiles of *sasA* expression, average OPP incorporation in the top two quartiles as compared to the bottom quartile is significantly higher (Fig 2D). In comparison, there was no significant difference in OPP incorporation between any of the quartiles of *sasB* expression (Fig 2C). Thus, differences in *sasA*, but not *sasB*, expression are associated with the observed heterogeneity in protein synthesis.

**Fig 2.**
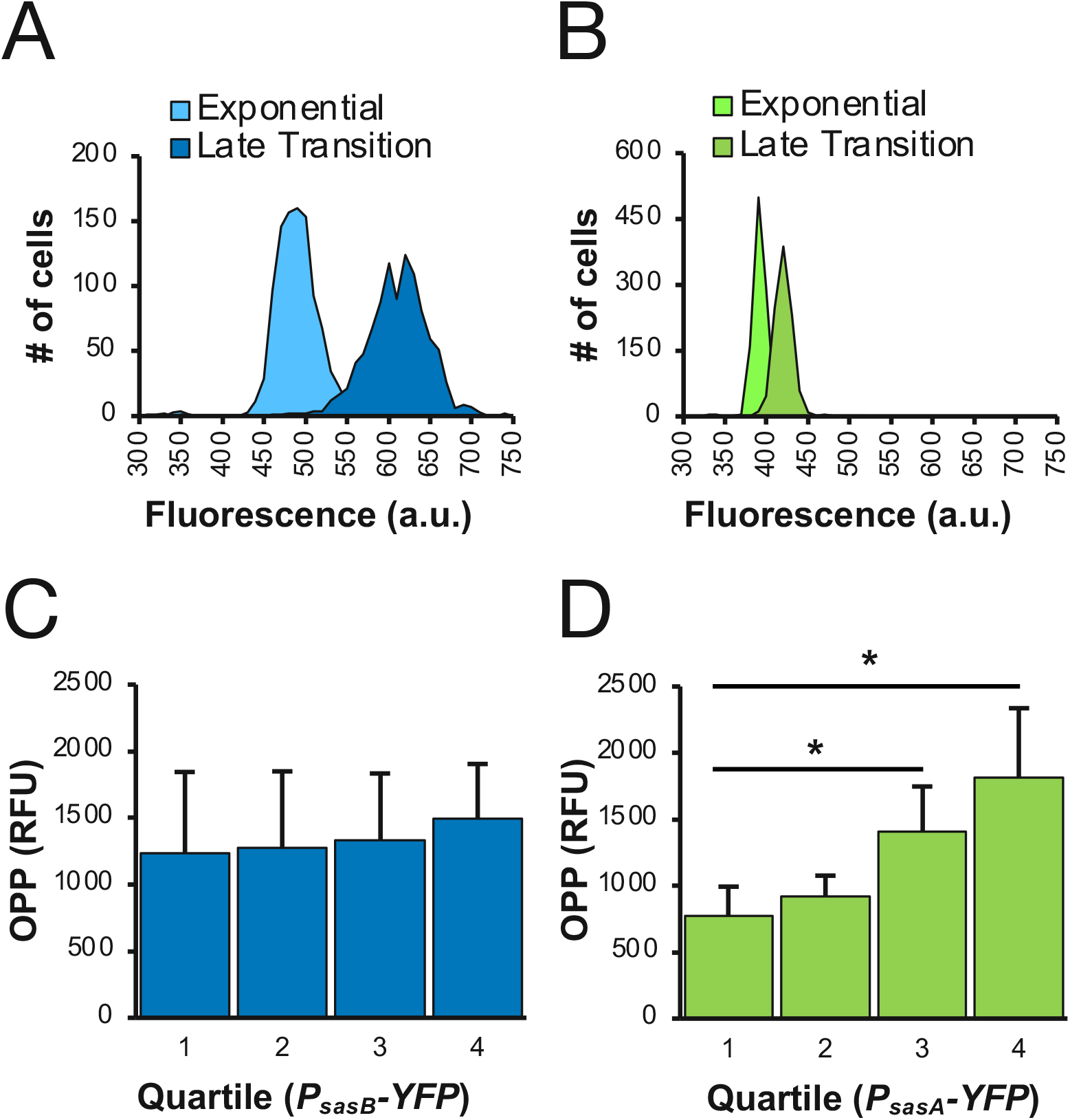
Relationship between *sasA* or *sasB* expression and OPP incorporation. **(A, B)** Representative population distribution of *B. subtilis* carrying a transcriptional reporter of **(A)** *P*_*sasB*_*-yfp* (JDB4341) or **(B)** *P*_*sasA*_*-yfp* (JDB4030) during exponential and late transition phase. **(C, D)** Average OPP incorporation of each quartile of **(C)** *P*_*sasB*_*-yfp* expression or **(D)** *P*_*sasA*_*-yfp* expression from lowest to highest. Statistical analysis (one tailed t-test) showed no significant difference in OPP incorporation between any *P*_*sasB*_*-YFP* quartiles (p>0.05) and significantly higher OPP incorporation between quartiles 1 and 3 and quartiles 1 and 4 of *P*_*sasA*_*-yfp* expression (p-values 0.027 and 0.016, respectively).

### SasB allosteric activation is necessary for heterogeneity

If changes in *sasB* transcription are not associated with differences in protein synthesis (Fig 2C), but SasB is necessary for the heterogeneity of protein synthesis (Fig 1B), what mechanism underlies differential SasB activity in single cells? *B. subtilis* SasB is subject to allosteric activation by pppGpp, the main product of *B. subtilis* RelA (14). Phe-42 is a key residue in this activation and a SasB mutant protein carrying an F42A substitution (SasB^F42^) is not allosterically activated by pppGpp *in vitro* (14). We investigated the importance of this allosteric activation for protein synthesis heterogeneity using a strain expressing SasB^F42^. Heterogeneity of this strain is significantly attenuated compared to the WT strain, demonstrating the importance of the allosteric activation of SasB by pppGpp for the bimodality of protein synthesis activity (Fig 3A, B; S3 Fig).

**Fig 3.**
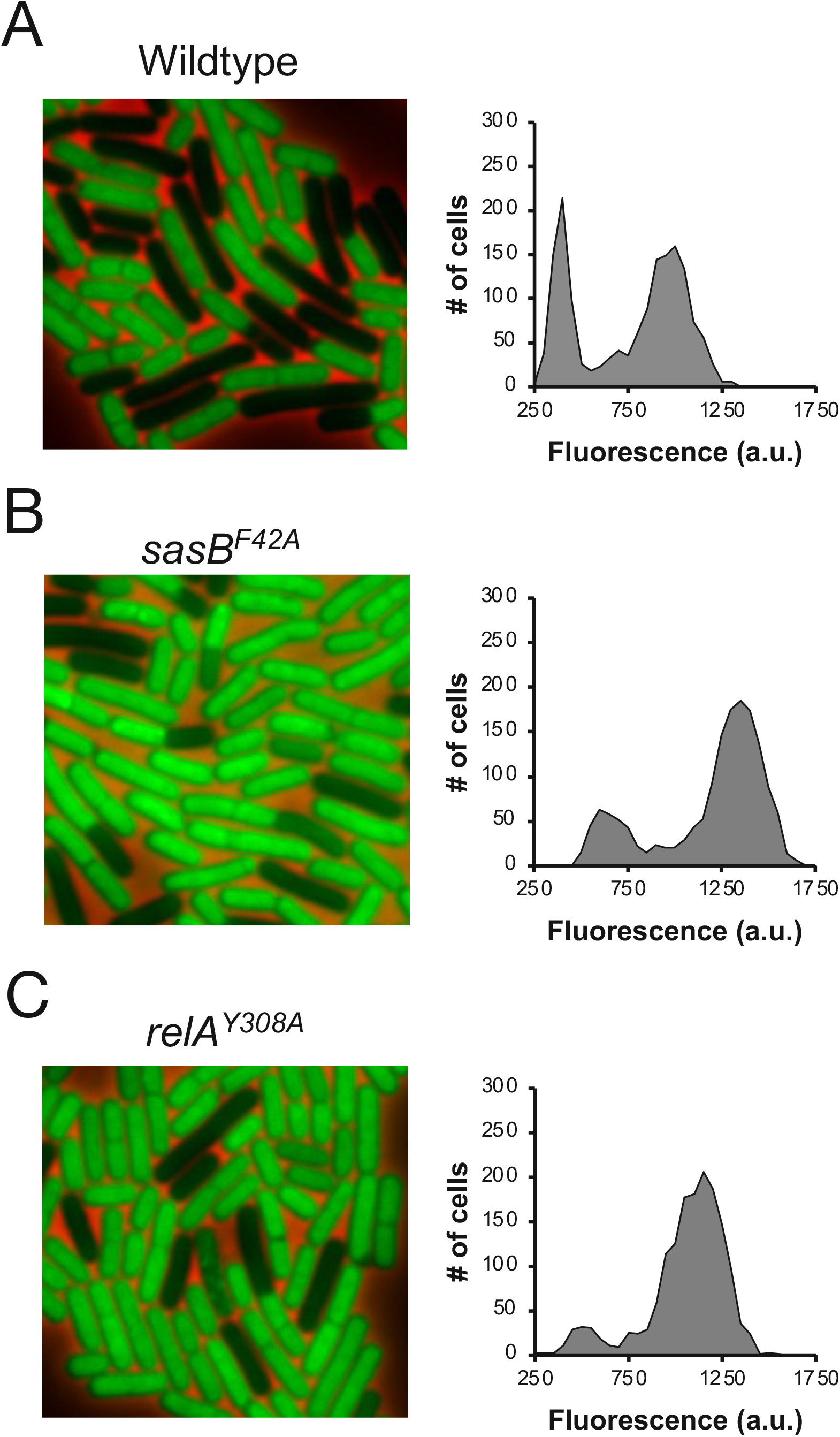
Allosteric activation of SasB is required for bimodality during exit from rapid growth. **(A, B,C)** Representative picture and population distribution of OPP labeled **(A)** wildtype (JDB1772), **(B)** *sasB*^*F42A*^ (JDB4340), and **(C)** *relA*^*Y308A*^ (JDB4300) strains during late transition phase.

This result suggests that the enzyme responsible for pppGpp synthesis could also affect the heterogeneity. RelA is the primary source of pppGpp in *B. subtilis* (11), so the loss of *relA* would be predicted to affect SasB activity. We therefore generated a strain expressing a RelA mutant protein (RelA^Y308A^) carrying a single amino acid change at a conserved residue essential for synthase but not hydrolysis activity (15, 16) since RelA hydrolytic activity is essential in a strain that retains functional *sasA* and *sasB* genes (17). Labeling of this strain with OPP in late transition phase revealed that the “OFF” population was largely absent (Fig 3C; S3), demonstrating that RelA-mediated pppGpp synthesis is important for the bimodality.

### SasB allosteric activation is inhibited by pGpp

A strain lacking SasA (Δ*sasA*) contains more “OFF” cells as compared to the wildtype parent (Fig 1C). The presence of this sub-population of cells depends on a SasB protein that can be allosterically activated (Fig 3B). Integrating these two observations, we hypothesized that the product of SasA (pGpp) inhibits the allosteric activation of SasB by pppGpp. The similarity of pGpp and pppGpp suggests that they could have an antagonistic interaction since they are likely capable of binding to the same site on SasB, but their differing phosphorylation states could affect their ability to allosterically activate SasB.

We tested this model by assaying *in vitro* whether pGpp inhibits the allosteric activation of SasB. First, we confirmed that SasB generates more ppGpp when reactions are supplemented with pppGpp (14) and observed a ∼2 fold increase in ppGpp production when SasB was incubated with pppGpp (Fig 4A). Using pGpp synthesized *in vitro* by the recently identified (p)ppGpp hydrolase NahA (18), we observed that pGpp attenuates the allosteric activation of SasB in a dose dependent manner (Fig 4A). Since even the highest concentration of pGpp did not decrease production of ppGpp relative to that generated by SasB without the addition of pppGpp (Fig 4A), the inhibition is likely specific to the allosteric activation. We first tested this directly by assaying the effect of pGpp on SasB activity in the absence of its allosteric activator (pppGpp). Addition of pGpp did not significantly affect SasB activity within the range of pGpp concentrations we used previously (Fig S4). We further confirmed the specificity by assaying a SasB^F42^ mutant protein that is insensitive to allosteric activation by pppGpp (14). As previously reported SasB^F42A^ has similar activity to a non-allosterically activated WT SasB in the presence of pppGpp (Fig 4B). However, in contrast with wildtype SasB, pGpp does not affect the activity of SasB^F42A^ even when pppGpp is included (Fig 4B).

**Fig 4.**
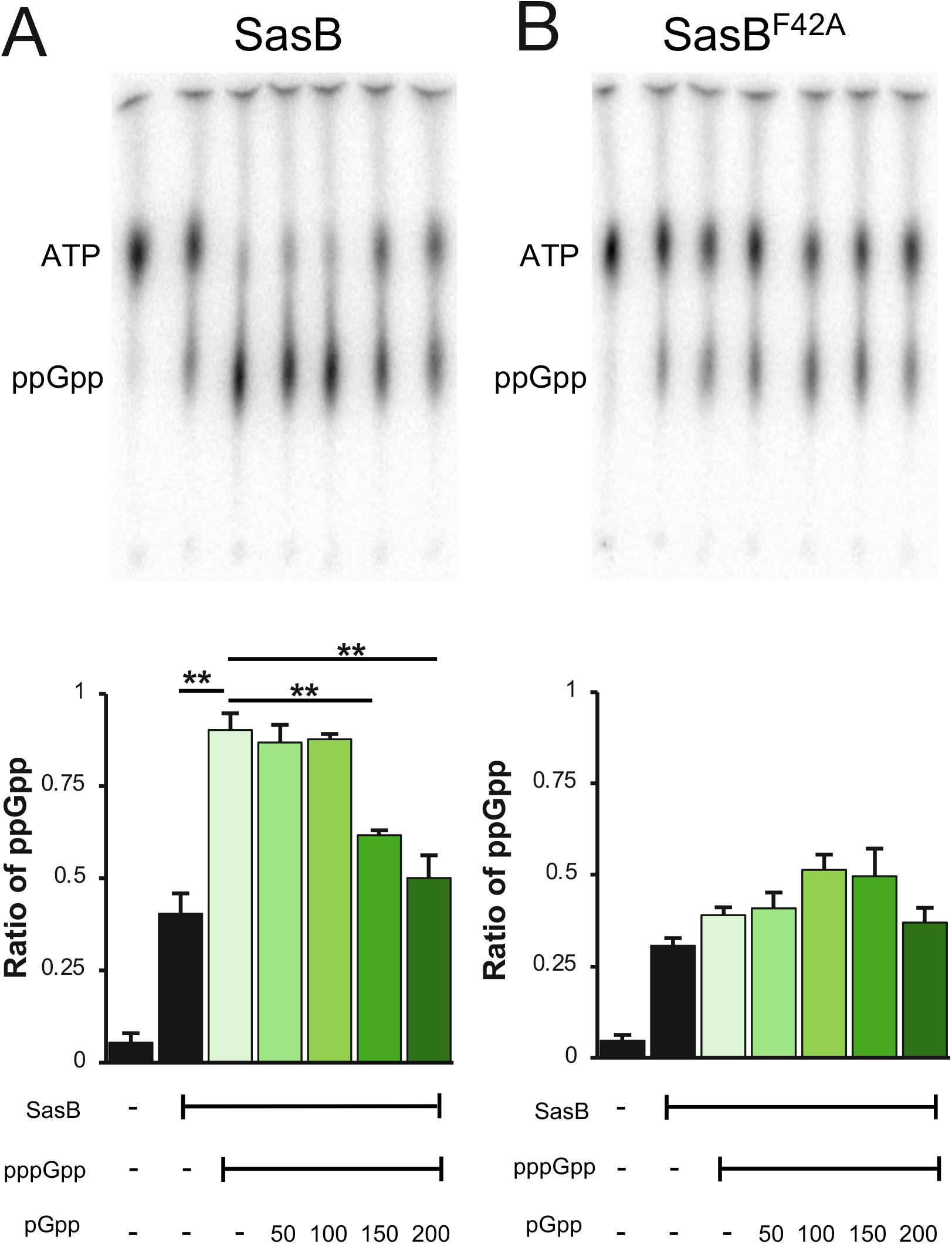
The product of SasA (pGpp) inhibits the allosteric activation of SasB by pppGpp. **(A)** Representative TLC of nucleotides present following incubation of wildtype SasB with [α-^32^P]-ATP and GDP in the presence or absence of pppGpp and increasing concentrations of pGpp (top). Quantitation of the ratio of ppGpp to total nucleotides present in each lane in TLC. Ratio of ppGpp was calculated using the formula: ppGpp/ATP + ppGpp (bottom) **(B)** Representative TLC of nucleotides present following incubation of SasB^F42A^ with [α-^32^P]-ATP and GDP in the presence or absence of pppGpp and increasing concentrations of pGpp (top). Ratio of ppGpp present in each lane in TLC as determined the formula, ppGpp/ATP + ppGpp (bottom). Statistical analysis (two tailed t-test) showed no significance (p > 0.05) between reactions containing SasB in the presence or absence of pppGpp and/or pGpp.

These *in vitro* biochemical experiments suggest that the effect of SasA on protein synthesis heterogeneity is dependent on the activity of SasB. If this is true *in vivo*, then the phenotype of a Δ*sasA* mutation should be epistatic to that of a Δ*sasB* mutation. Consistently, the population of “OFF” cells in a Δ*sasA* strain is absent in a strain lacking both SasA and SasB (*sasA sasB*) (Fig 5A, B; S5). Thus, the effect of SasA is dependent *in vivo* on SasB. Finally, since RelA activates SasB, a Δ*sasA* mutation should be epistatic to a *relA* mutation with respect to protein synthesis. A strain expressing RelA^Y308A^ and carrying a Δ*sasA* mutation exhibits a loss of heterogeneity similar to the *relA*^*Y308A*^ strain, demonstrating that the effect of the Δ*sasA* mutation depends on a functional RelA synthase (Fig 5A, C; S5). This result is consistent with the hypothesis that *sasA* is epistatic to *relA*.

**Fig 5.**
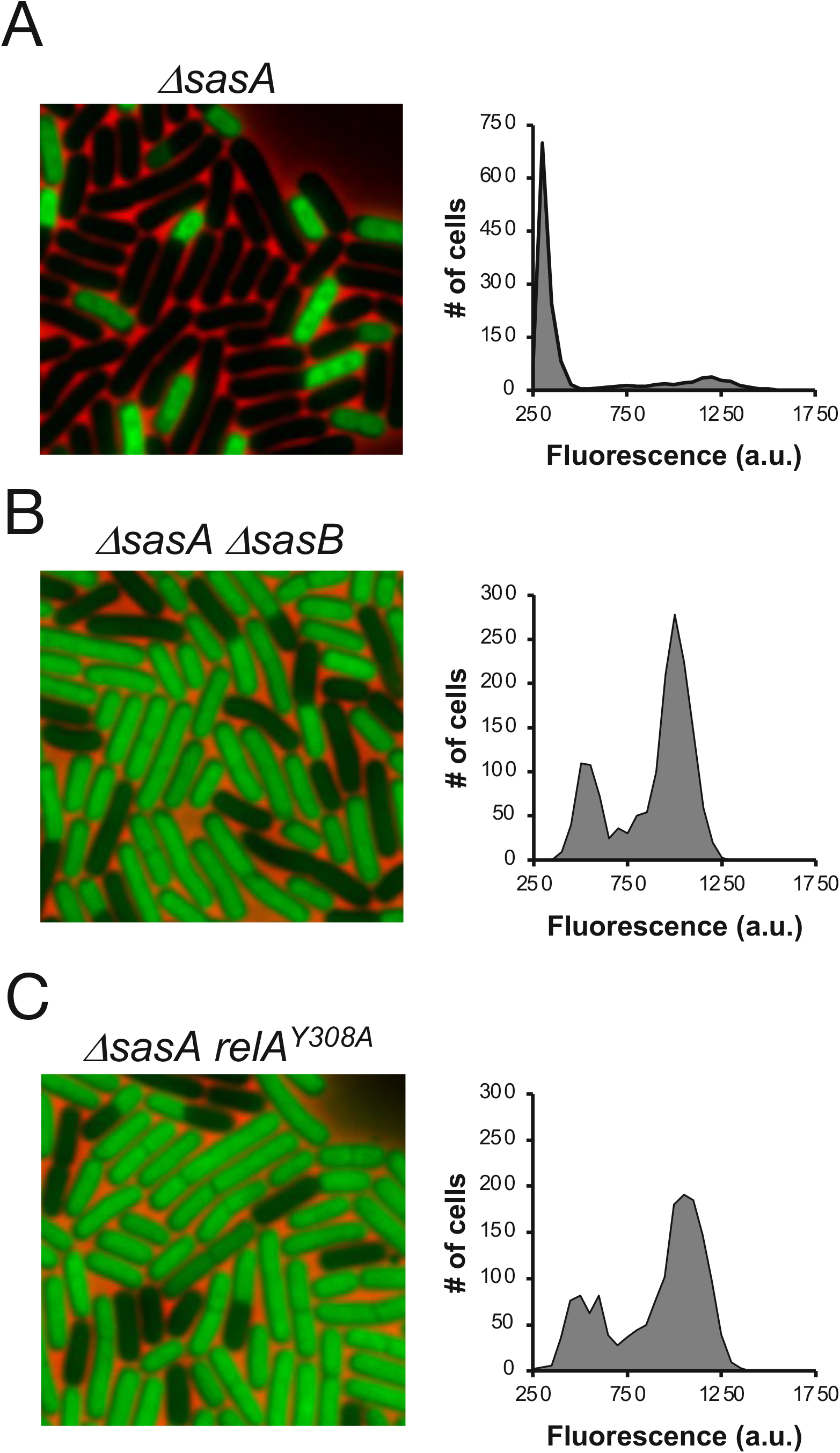
*sasA* is epistatic to *sasB* and *relA*. **(A, B, C)** Representative pictures and population distributions of OPP labeled **(A)** *ΔsasA* (JDB4310), **(B)** *ΔsasA ΔsasB* (JDB4312) **(C)** Δ*sasA relA*^*Y308A*^ (JDB 4301) strains during late transition phase.

While SasA is the only known (pp)pGpp synthase that predominately produces pGpp *in vivo* in *B. subtilis* (12), pGpp also accumulates in stationary phase cells as a result of degradation of both ppGpp and pppGpp by the (p)ppGpp hydrolase NahA (18, 19). We therefore asked if NahA contributes to the heterogeneity in protein synthesis by comparing OPP incorporation in wildtype and Δ *nahA* cells during late transition phase. We observed no difference in heterogeneity (Fig S6) consistent with SasA being the primary source of pGpp.

## Discussion

*B. subtilis* populations experiencing nutrient limitation and entering into quiescence respond bimodally with respect to global protein synthesis activity (5). Here, we find that this bimodality depends on all three (pp)pGpp synthases. We demonstrate that it is dependent on the allosteric activation of SasB by the RelA product pppGpp and that this activation is antagonized by the SasA product pGpp (Fig 6A). Our work therefore provides a mechanism for the phenotypic heterogeneity observed and identifies novel regulatory interactions between (pp)pGpp synthases.

**Fig 6.**
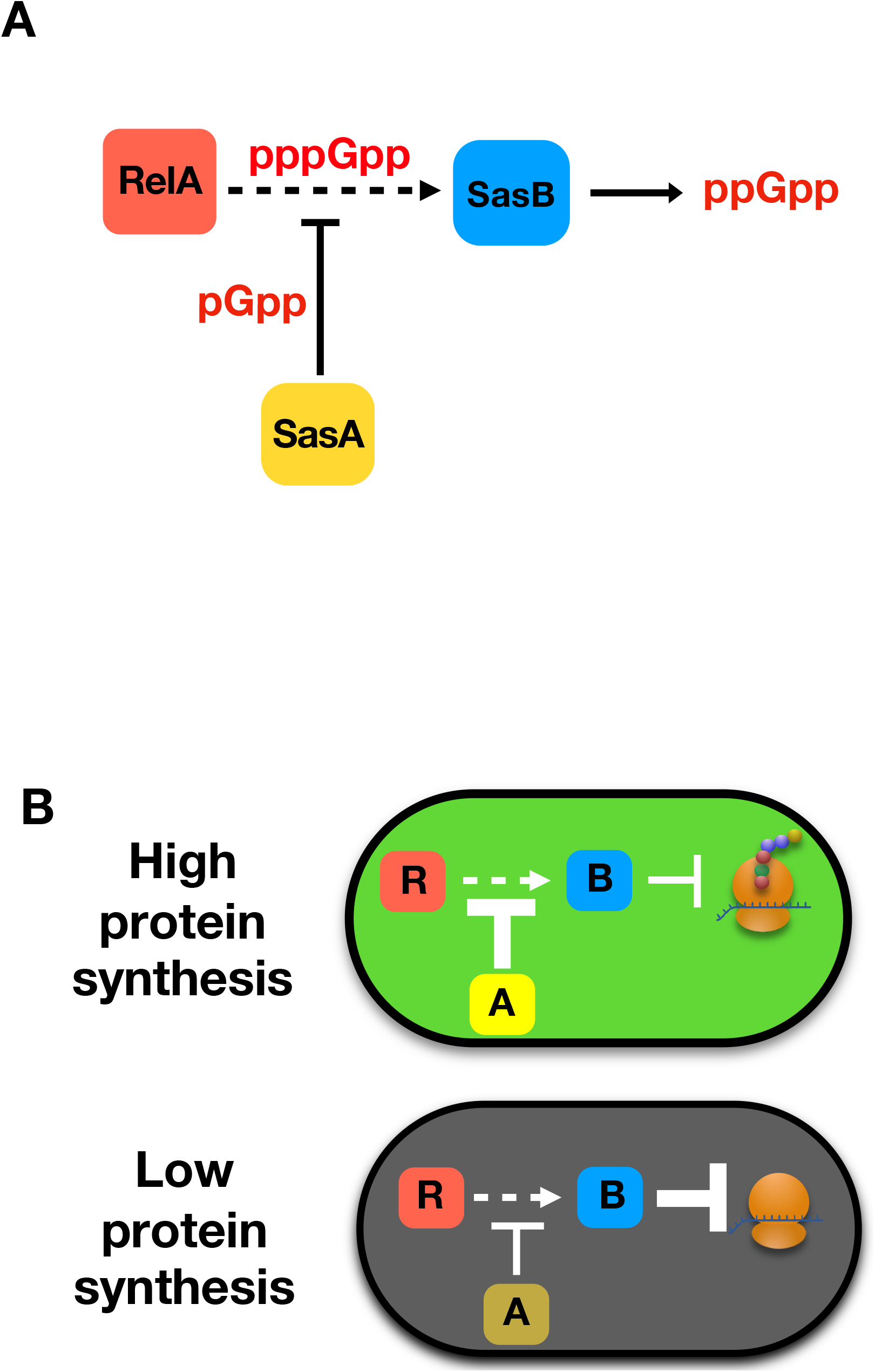
Generation of heterogeneity in protein synthesis. **(A)** In response to amino acid limitation, RelA synthesizes pppGpp that allosterically activates SasB. This activation is inhibited by pGpp, the product of SasA and the crosstalk of these two nucleotides determines how much ppGpp SasB produces. **(B)** In cells with relatively higher *sasA* (‘A’) expression, increased inhibition of SasB (‘B’) allosteric activation by RelA (‘R’) results in relatively high protein synthesis. In cells with relatively lower *sasA* expression, decreased inhibition of SasB allosteric activation attenuates protein synthesis to a greater extent than in cells with lower *sasA* expression.

### Regulation of protein synthesis during nutrient limitation

The downregulation of protein synthesis in *B. subtilis* cells experiencing nutrient limitation occurs as a result of ppGpp binding and thereby inhibiting IF2 (5). SasB is the main source of ppGpp and this work identifies how ppGpp synthesis by SasB and the subsequent downregulation of protein synthesis is coupled to changes in environmental conditions. First, SasB allosteric activation by the RelA product pppGpp is required for the downregulation of protein synthesis in a subpopulation of cells (Figs 3, 5). RelA activity reflects tRNA charging levels (11), thereby coupling SasB-dependent regulation of protein synthesis to amino acid availability. Second, the SasA product pGpp inhibits the allosteric activation of SasB (Fig 4). Although SasA is constitutively active, *sasA* expression, at least in part, reflects availability of the Lipid II peptidoglycan precursor (20-22), thereby coupling SasB-dependent regulation of protein synthesis to cell wall metabolism. Thus, the roles of RelA and SasA in regulating SasB activation provides a mechanism to integrate multiple environmental signals in the decision to attenuate protein synthesis.

### Physiological sources of variability in SasB activity

Phenotypic heterogeneity such as that observed here in the context of protein synthesis can arise from stochastic differences in gene expression (23). Although *sasB* expression exhibits substantial variability in expression cell to cell (Fig 2A), it does not correspond with the level of protein synthesis in individual cells (Fig 2C). Thus, variability of SasB activity in single cells is likely relevant. What could be responsible? Our observations link heterogeneity to the convergent regulation of SasB allosteric activation by the products of the RelA and SasA synthases, pppGpp and pGpp, respectively (Fig 4A). Thus, both enzymes are potential sources of variability and, consistently, strains carrying either *relA*^*Y308A*^ or Δ*sasA* mutations exhibit a loss in heterogeneity as compared to the wildtype (Figs 1C, 3C). Since RelA is a cellular sensor of tRNA charging, levels of which are highly sensitive to growth conditions (24), variations in this parameter could contribute to variability in protein synthesis via modulation of RelA activity. Noise in *sasA* transcription is dependent on the activity of PrkC, a membrane Ser/Thr kinase that regulates *sasA* via the essential WalRK two component system (20). Since both WalRK (25) and PrkC (26) activities reflect cell wall metabolism, variation in this process could also impact *sasA* variability. Thus, differences in the protein synthesis activity of individual cells may reflect cellular variations in amino acid and cell wall metabolism.

### Allosteric activation of (pp)pGpp synthases

Many genes encoding SAS proteins such as *sasB* are transcriptionally regulated (9). In addition, we observe here there that allosteric activation of SasB by pppGpp (14) is required for the attenuation of protein synthesis (Fig 3) demonstrating *sasB* transcription is necessary but not sufficient, at least in the physiological context of nutrient limitation. We also find that this allosteric activation is antagonized by the SasA product pGpp (Fig 6A), consistent with the epistatic relationship between *sasB* and *sasA* (Fig 5A). Antagonistic regulatory mechanisms are likely widespread in this family of synthases. For example, the SasB homolog *Enterococcus faecalis* RelQ is attenuated by RNA that competes with pppGpp for binding to the allosteric site (27). Given the very recently observed allosteric activation of *B. subtilis* RelA by (p)ppGpp (28), an important question for future study is to determine whether this activation is also subject to antagonism by pGpp and, if so, to characterize the physiological consequences of this regulation.

### (pp)pGpp synthases

The different protein synthesis activity of strains carrying a mutation in one of the genes encoding a (pp)pGpp synthase (Figs 1, 3C) is consistent with previous reports that SAS enzymes differ between themselves and also with RelA in the guanosine nucleotide that they preferentially produce (18, 29-31). Our observations demonstrate that each particular product differs in its *in vivo* function, thereby extending previous observations that ppGpp and pppGpp can differ in their effect on gene transcription in *E. coli* (32). The biochemical experiments demonstrating that pGpp antagonizes pppGpp allosteric activation of SasB, but itself is not capable of activation (Fig 4A; S4) are consistent with our physiological experiments. The biochemical activity of these nucleotides have been reported to differ, including observations that pppGpp is much more potent than ppGpp in stimulating SasB (14), that pGpp is a significantly more potent inhibitor of purine salvage enzyme XPRT than ppGpp (33), and that ppGpp, but not pppGpp, inhibits the function of IF2 in stimulating subunit joining (34). Thus, these three closely related nucleotides have distinct biochemical and, as we show here, physiological activities.

### Physiological implications of heterogeneity in protein synthesis

(p)ppGpp has long been thought to mediate entry into bacterial quiescence (35, 36). This transition facilitates survival in nutrient limited environments and its regulation depends upon the integration of a multitude of rapidly changing environmental signals that themselves may impair decision-making. One way bacteria deal with such uncertainty is to generate subpopulations, with distinct, often bimodal phenotypes from a population of genetically identical cells (23). Examples of phenotypic variation in *B. subtilis* include heterogeneity in specific metabolic activities such as acetate production (37) or in developmental transitions such as sporulation (38) and competence (39). The phenotypic variation in protein synthesis activity we observe here has potentially broad functional implications given its central role in cellular physiology. A global reduction in protein synthesis activity, if accompanied by a constant rate of protein degradation, would have the effect of reducing overall metabolic capacity, especially by affecting processes like ribosome assembly. Global effects also could have specific regulatory consequences. For example, the alternative sigma factor *B. subtilis* SigD drives expression of genes controlling daughter cell separation and motility that exhibit well characterized phenotypic variation. RelA affects both this variability as well as absolute levels of SigD (40), suggesting that differences in protein synthesis between cells contribute to SigD variability.

In summary, this work demonstrates that three differentially phosphorylated nucleotides and their respective synthases comprise a signaling network responsible for the heterogenous regulation of protein synthesis as *B. subtilis* cultures enter quiescence. We find that this heterogeneity is dependent on the RelA product pppGpp, which allosterically activates SasB, and the SasA product pGpp, which antagonizes this activation. Since the synthesis of pppGpp and pGpp reflects amino acid and peptidoglycan precursor availability, respectively, these parameters are thereby coupled to protein synthesis activity and facilitate cell decision making during the entry into quiescence.

## Materials and Methods

### Strains and media

Strains were derived from *B. subtilis* 168 *trpC2. sasA* (*ywaC*) and *sasB* (*yjbM*) gene knockouts were from transformed into *B. subtilis* 168 *trpC2* using genomic DNA from BD5467 (41). The *sasB* transcriptional reporter strain was constructed similarly as described (20). Briefly, a 107 bp region encompassing the *sasB* operon promoter (*P*_*sasB*_) was amplified and inserted into AEC 127 using *EcoR*I and *BamH*I sites. The resulting AEC 127 *P*_*sasB*_ was integrated into *B. subtilis* 168 *trpC2* at *sacA. sasB*^*F42A*^ and *relA*^Y308A^ strains were generated using integration of pMINIMAD2 derivatives (pMINIMAD2 *sasB*^*F42A*^ and pMINIMAD2 *relA*^Y308A^, respectively). Briefly, *sasB* was amplified excluding start and stop codons and F42A mutation was introduced using overlap extension PCR. *sasB*^*F42A*^ was inserted into pMINIMAD2 vector using *EcoR*I and *Sal*I sites. pMINIMAD2 *sasB*^*F42A*^ vector was transformed into *B. subtilis* 168 *trpC2* using a standard transformation protocol. Transformants were selected for erythromycin resistance at 45 °C overnight and grown for 8 hours at RT in LB. Cultures were diluted 1:10 in LB and grown overnight. Cultures were plated for single colonies and grown overnight at 45 °C. Single colonies were checked for erythromycin sensitivity and sensitive clones were checked for *sasB*^*F42A*^ allele by Sanger sequencing of *sasB* amplified genomic region. The *relA*^Y308A^ strain was generated in a similar way but *EcoR*I and *BamH*I sites were used to insert the *relA*^Y308A^ gene into pMINIMAD2.

### Growth curves

Growth curves were performed in a Tecan Infinite m200 plate reader at 37 °C with continuous shaking and OD_600_ measurements were made every five min. Cultures were grown from single colonies from fresh LB plates grown overnight at 37 °C. Exponential phase starter cultures (OD_600_∼ 0.5 - 1.5) were diluted to OD_600_ = 0.01 and grown in 96-well Nunclon Delta surface clear plates (Thermo Scientific) with 150 μL per well. All growth curves were done in triplicate and media-only wells were used to subtract background absorbance.

### OPP labeling

OPP labeling of cells was as described (5). Exposure times were 30 msec for phase contrast, and 20 msec for mCherry. Fluorescence intensity of ∼1570 single cells per experiment was determined using ImageJ. Cells were binned based on fluorescence intensity using 50 a.u. wide bins in all experiments and number of cells in each bin presented as a histogram.

### Protein expression and purification

Wildtype and F42A SasB proteins were expressed and purified essentially as described (14). Wildtype *sasB* was amplified from *B. subtilis* 168 *trpC2*. The F42A mutation was introduced using overlap extension PCR. WT and *sasB*^*F42A*^ PCR products were inserted into pETPHOS expression vector using *EcoR*I and *BamH*I sites. pETPHOS WT *sasB* and pETPHOS *sasB*^*F42A*^ were transformed into *E*.*coli* BL21 and proteins were induced with 1 mM IPTG for 2h at OD_600_ ∼0.5.

Cells were harvested at 4 °C and lysed using a Fastprep (MP biomedicals) in 50 mM Tris (pH 8.0), 250 mM NaCl, 5 mM MgCl_2_, 2 mM BME, 0.2 mM PMSF, and 10mM imidazole. Lysates were clarified and bound to a Ni-NTA column (Qiagen) for 1h. Columns were washed using 20 mM imidazole. Protein was eluted using 500 mM imidazole, dialyzed into 20mM Tris, 500 mM NaCl, 5mM MgCl_2_, 2 mM BME, and 10% glycerol and stored at -20 °C. NahA protein was purified in a similar way except that NahA was induced for 1h at 30 °C and NahA expressing cells (JDE3138) were lysed, washed, and eluted in 250 mM NaCl instead of 500 mM.

### pGpp synthesis

pGpp was synthesized *in vitro* by purified NahA enzyme as described (18). Briefly, 10 nM purified *B. subtilis* NahA was incubated with 30 nM pppGpp (Trilink Biotechnologies) in 40 mM Tris-HCl (pH 7.5), 100 mM NaCl, 10 mM MgCl_2_ at 37 °C for 1 hour. Reactions were monitored for conversion of pppGpp to pGpp using thin layer chromatography on PEI-cellulose plates in 1.5 M KH_2_PO (pH 3.6). Nucleotides were visualized using short wave UV light. NahA enzyme was precipitated using ice cold acetone and nucleotides were stored at -20 °C.

### SasB activity assays and TLC

SasB activity was assayed by measuring the amount of ppGpp generated similar to (5). Briefly, 0.8 μM purified *B. subtilis* WT or F42A SasB was incubated with 0.5 μCi of [γ-^32^P]-ATP (PerkinElmer) and 50 μM GDP in 20 mM Tris (pH 7.5), 500 mM NaCl, 5 mM MgCl_2_, 2mM BME. SasB was allosterically activated using 12.5 μM pppGpp (Trilink Biotechnologies) and pGpp was added as noted. Reactions were performed in a total volume of 10 μL, and each reaction was incubated at 37 °C for 1 min before being stopped using 5 μL of ice cold acetone. Conversion of ATP to ppGpp was visualized using thin layer chromatography on PEI-cellulose plates in 1.5 M KH_2_PO_4_ (pH 3.6). Plates were dried completely at RT and exposed for 5 min on a phosphor storage screen and visualized (GE Typhoon). ATP and ppGpp spot intensities were quantified using ImageJ.

## Acknowledgements

SD was supported in part by the Columbia University Graduate Training Program in Microbiology, Immunology and Infection (R01 AI106711, Program Directors D. Fidock and L. Symington). JD was supported by NIH R01GM141953, R35GM141953, R21AI156397, and is a Burroughs-Welcome Investigator in the Pathogenesis of Infectious Disease.

## Supporting information

**S1 Fig.**
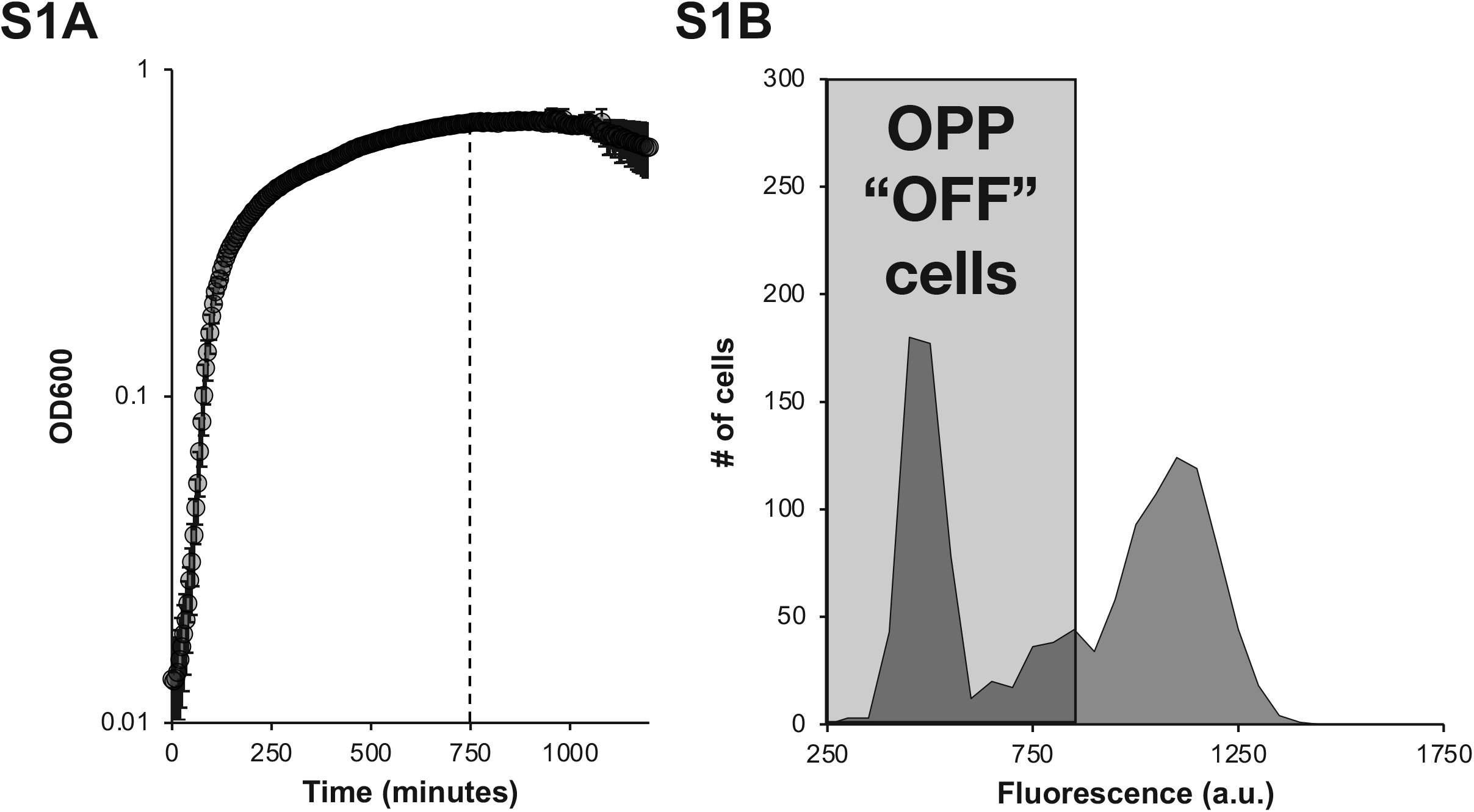
Late transition phase time point and determination of “OFF” cells. **(A)** Growth curve of wildtype *B. subtilis* showing point (OD_600_ ∼0.685) where cells were labeled with OPP (dashed line). **(B)** Representative distribution of OPP labeled wildtype *B. subtilis*. Gray box shows cutoff for cells with low rates of protein synthesis (“OFF”). Threshold was determined as the value (850 a.u.) that is higher than >95% of cells of wildtype *B. subtilis* that was labeled with OPP in stationary phase across three independent experiments.

**S2 Fig.**
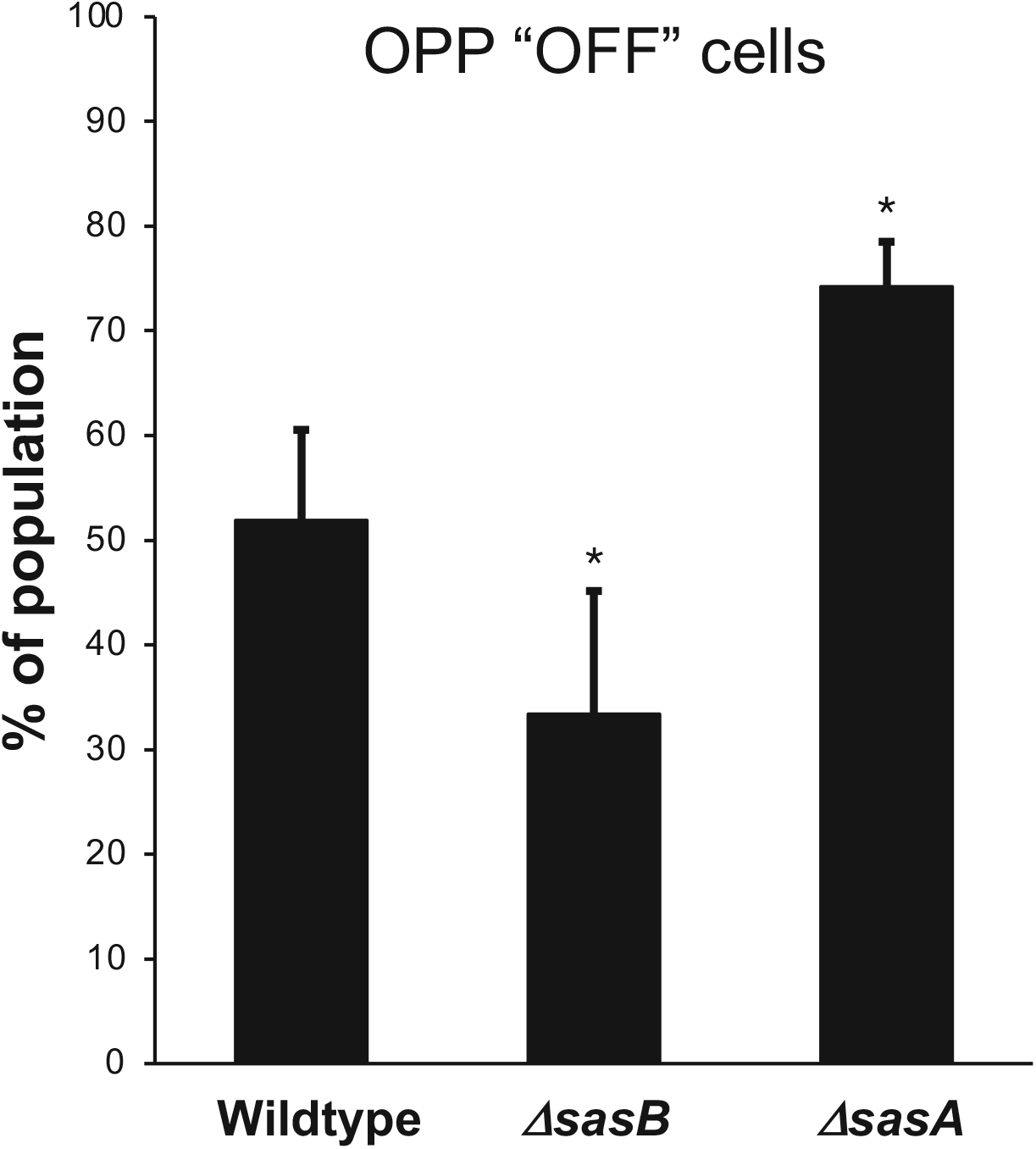
Quantitation of “OFF” cells and statistical analysis for Fig 1. Cells were designated as “OFF” using threshold in Fig S1. % “OFF” in a population was quantified in three separate experiments containing a total of> 1300 cells (means ± SDs). ^*^p < 0.05,

**S3 Fig.**
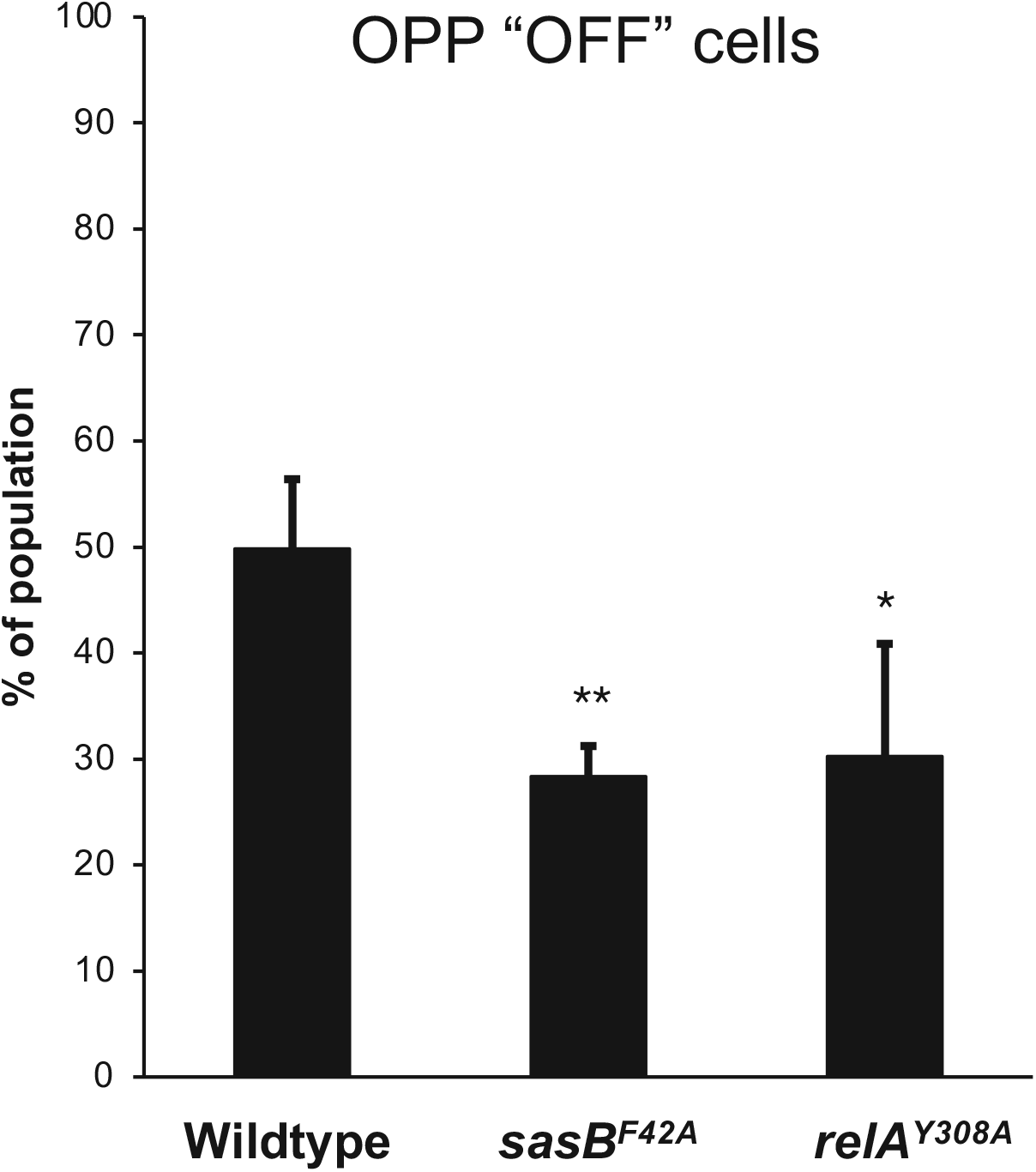
Quantitation of “OFF” cells and statistical analysis for Fig 3. Cells were designated as “OFF” using threshold in Fig S1. % of population “OFF” was quantified in three separate experiments containing a total of >1300 cells (means ± SDs). ^*^p < 0.05, ^**^p < 0.01.

**S4 Fig.**
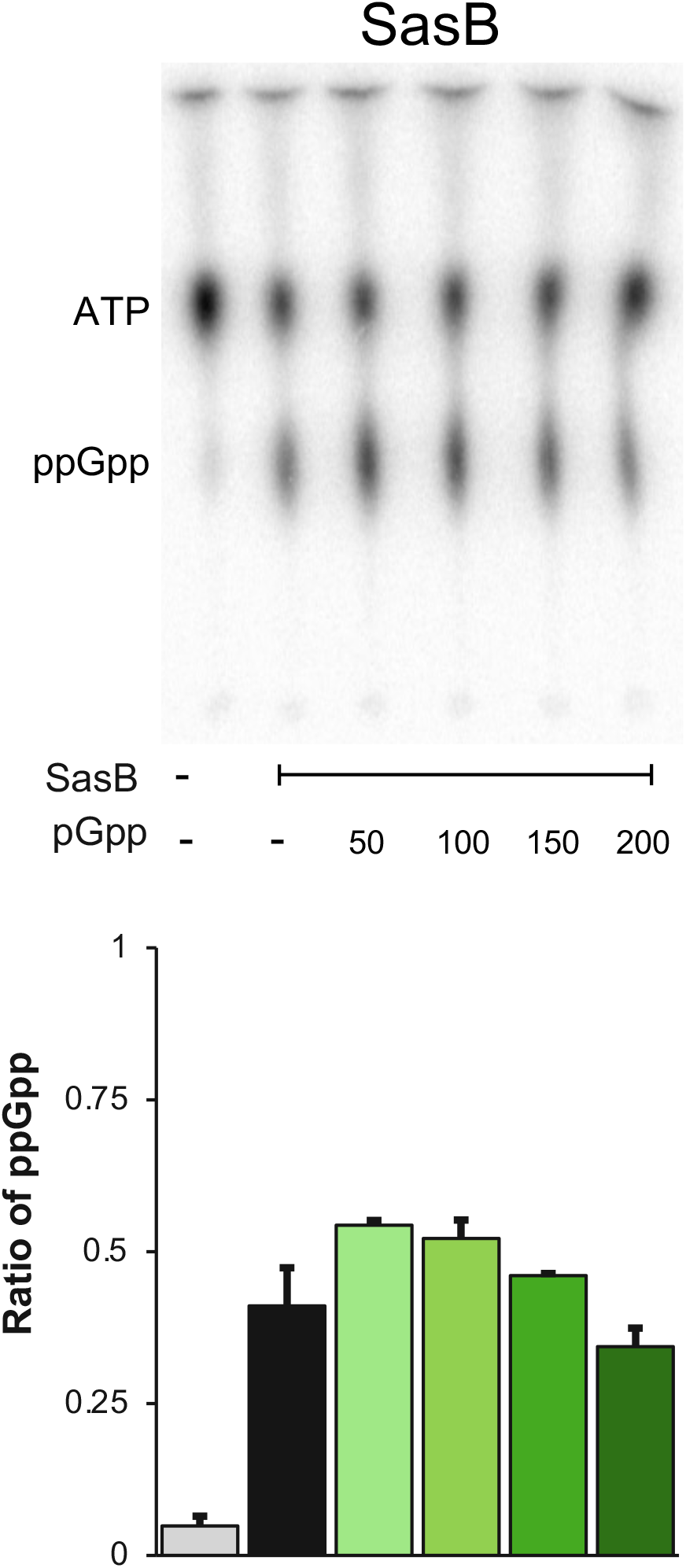
pGpp does not inhibit SasB basal activity. **(Top)** representative TLC analysis of wildtype SasB activity in the absence of allosteric activation (no pppGpp added) and with increasing concentrations of pGpp (uM). **(Bottom)** ratio of ppGpp calculated using the formula, ppGpp/ATP + ppGpp. Statistical analysis (t-test) showed no significance (p > 0.05) between any reaction containing SasB whether or not pGpp was included. Statistical analysis was performed on three separate experiments (means ± SDs).

**S5 Fig.**
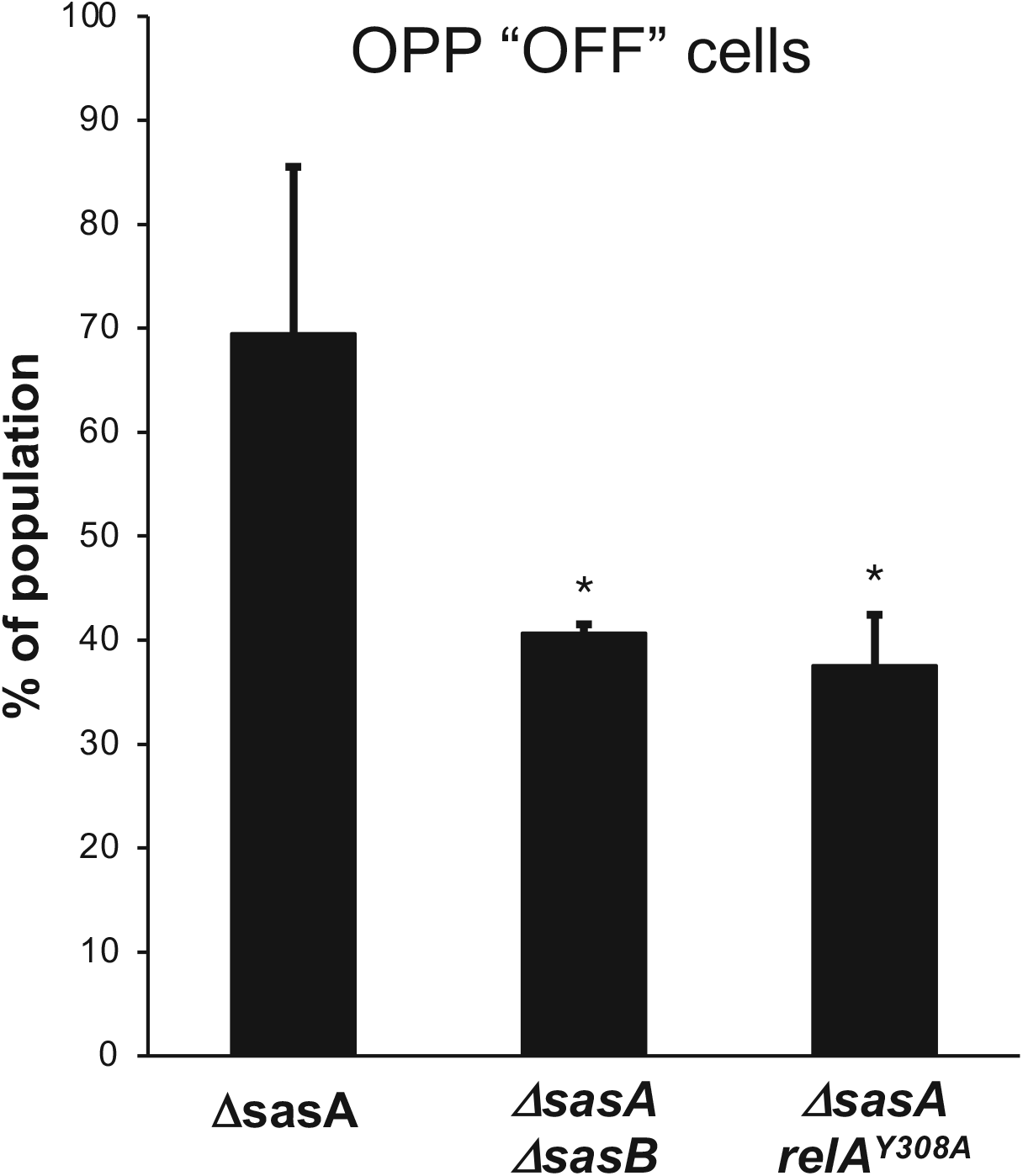
Quantitation of “OFF” cells and statistical analysis for Fig 5. Cells were designated as “OFF” using cutoff in Fig S1. % of population “OFF” was quantified in three separate experiments containing a total of >1300 cells (means ± SDs). ^*^p < 0.05

**S6 Fig.**
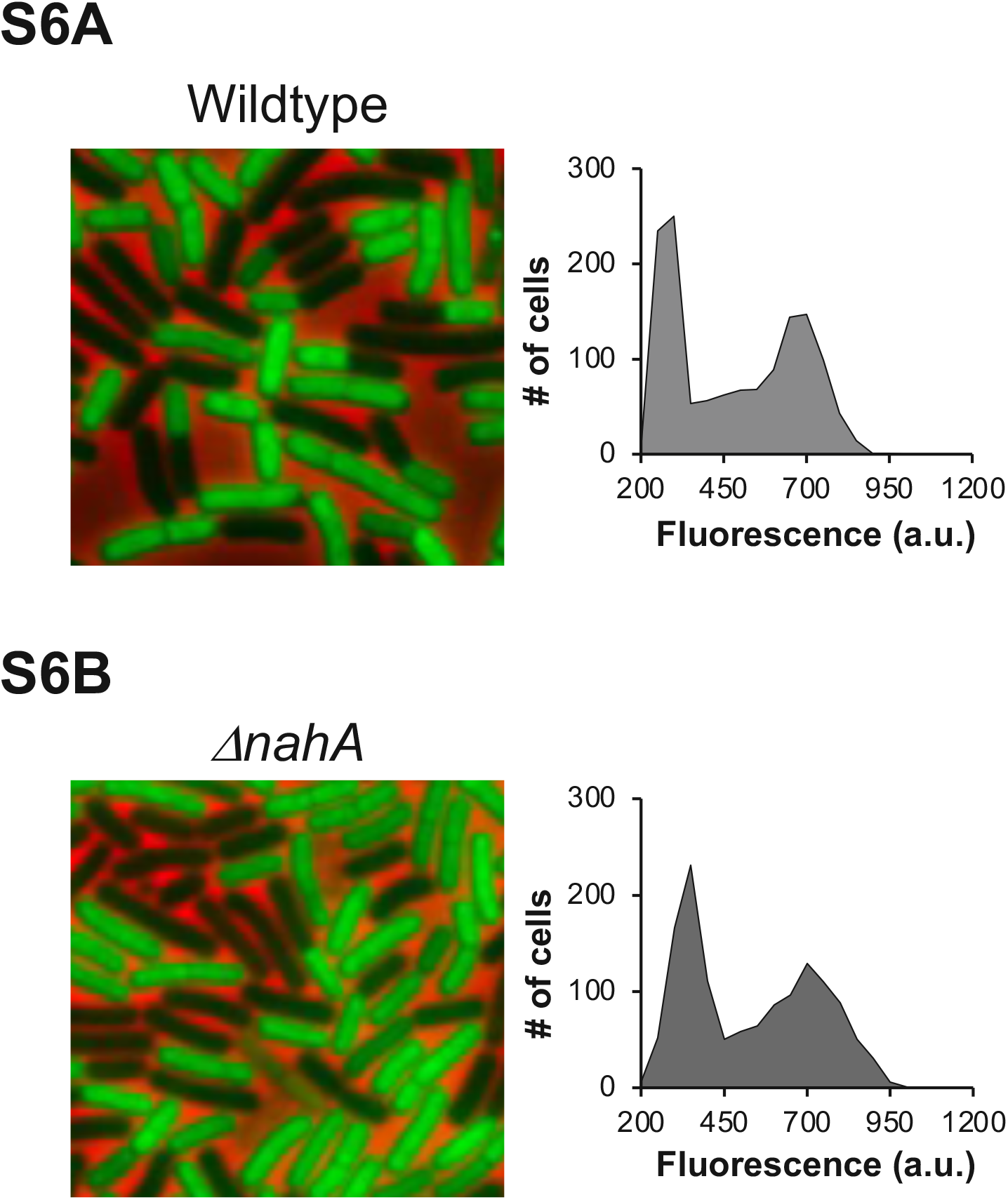
Effect of *nahA* on heterogeneity. **(A, B)** Representative picture and population distribution of OPP labeled **(A)** wildtype (JDB1772), **(B)** *ΔnahA* (JDB4095) strains during late transition phase.

**S1 Table.** Plasmids used in this study.

|  |  |
| --- | --- |
| pETPHOS | Lab stock |
| pMINIMAD2 | (6) |
| AEC 127 | (2) |
| pETPHOS WT *sasB* | This study |
| pETPHOS *sasB*<sup>F42A</sup> | This study |
| pETPHOS *yvcI* | This study |
| pMINIMAD2 *relA*<sup>Y308A</sup> | This study |
| pMINIMAD2 *sasB*<sup>F42A</sup> | This study |
| AEC 127 *PsasB* | This study |

**S2 Table.**
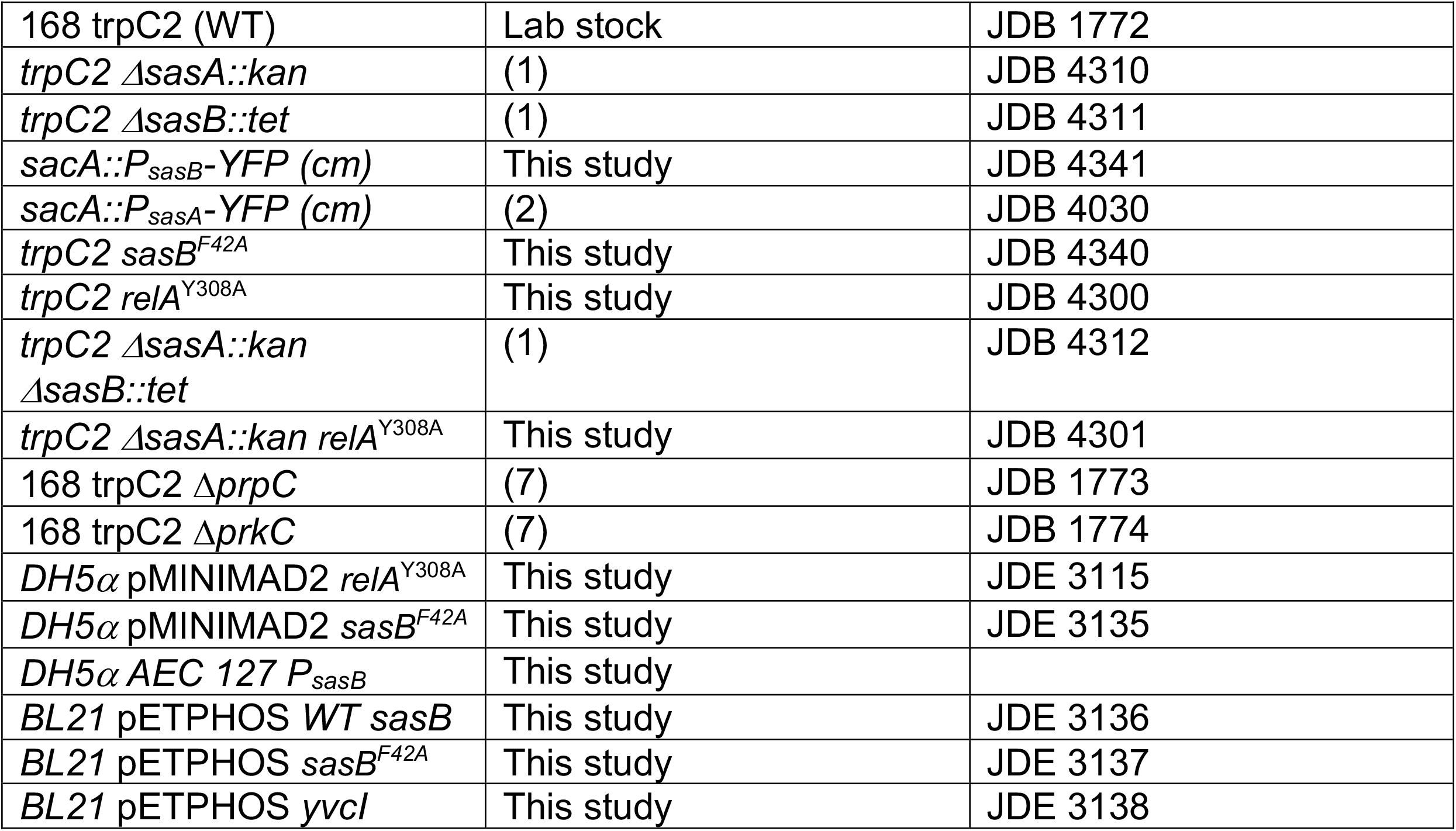
Strains used in this study.

**S3 Table.**
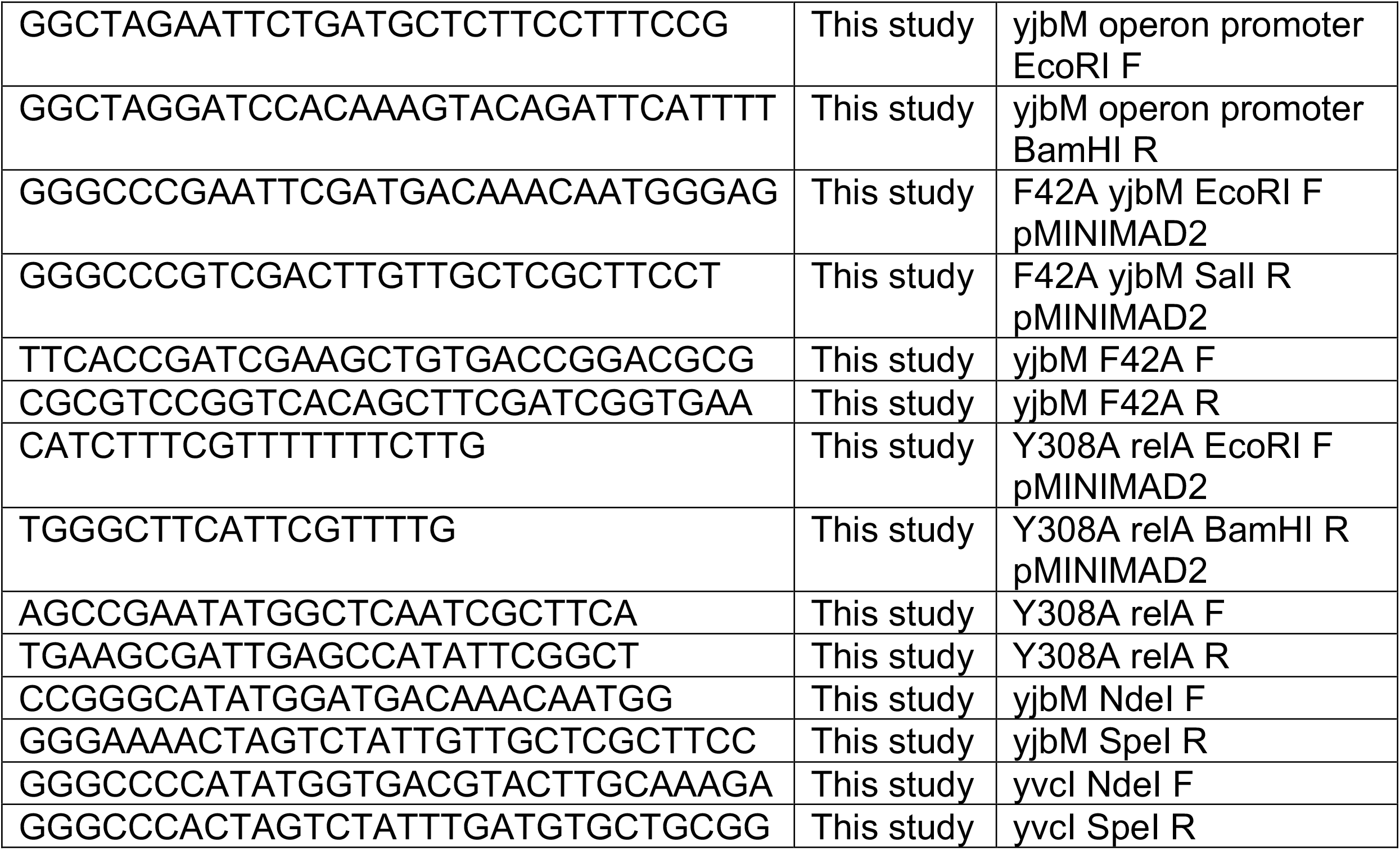
Oligonucleotides used in this study.

## Notes

### Competing Interest Statement

The authors have declared no competing interest.

## References

1. Gaca AO, Colomer-Winter C, Lemos JA. Many means to a common end: the intricacies of (p)ppGpp metabolism and its control of bacterial homeostasis. J Bacteriol. 2015;197(7):1146–56.

2. Steinchen W, Bange G. The magic dance of the alarmones (p)ppGpp. Mol Microbiol. 2016;101(4):531–44.

3. Artsimovitch I, Patlan V, Sekine S, Vassylyeva MN, Hosaka T, Ochi K, et al. Structural Basis for Transcription Regulation by Alarmone ppGpp. Cell. 2004;117:299–310.

4. Ross W, Sanchez-Vazquez P, Chen AY, Lee JH, Burgos HL, Gourse RL. ppGpp Binding to a Site at the RNAP-DksA Interface Accounts for Its Dramatic Effects on Transcription Initiation during the Stringent Response. Mol Cell. 2016;62(6):811–23.

5. Diez S, Ryu J, Caban K, Gonzalez RL, Jr., Dworkin J. The alarmones (p)ppGpp directly regulate translation initiation during entry into quiescence. Proc Natl Acad Sci U S A. 2020;117(27):15565– 72.

6. Kriel A, Bittner AN, Kim SH, Liu K, Tehranchi AK, Zou WY, et al. Direct regulation of GTP homeostasis by (p)ppGpp: a critical component of viability and stress resistance. Mol Cell. 2012;48(2):231–41.

7. Wang JD, Sanders GM, Grossman AD. Nutritional control of elongation of DNA replication by (p)ppGpp. Cell. 2007;128(5):865–75.

8. Corrigan RM, Bellows LE, Wood A, Grundling A. ppGpp negatively impacts ribosome assembly affecting growth and antimicrobial tolerance in Gram-positive bacteria. Proc Natl Acad Sci U S A. 2016;113(12):E1710–9.

9. Irving SE, Corrigan RM. Triggering the stringent response: signals responsible for activating (p)ppGpp synthesis in bacteria. Microbiology. 2018;164(3):268–76.

10. Steinchen W, Vogt MS, Altegoer F, Giammarinaro PI, Horvatek P, Wolz C, et al. Structural and mechanistic divergence of the small (p)ppGpp synthetases RelP and RelQ. Sci Rep. 2018;8(1):2195.

11. Wendrich TM, Marahiel MA. Cloning and characterization of a relA/spoT homologue from Bacillus subtilis Molecular Microbiology. 1997;26(1):65–79.

12. Tagami K, Nanamiya H, Kazo Y, Maehashi M, Suzuki S, Namba E, et al. Expression of a small (p)ppGpp synthetase, YwaC, in the (p)ppGpp(0) mutant of Bacillus subtilis triggers YvyD-dependent dimerization of ribosome. Microbiologyopen. 2012;1(2):115–34.

13. Nanamiya H, Kasai K, Nozawa A, Yun CS, Narisawa T, Murakami K, et al. Identification and functional analysis of novel (p)ppGpp synthetase genes in Bacillus subtilis. Mol Microbiol. 2008;67(2):291–304.

14. Steinchen W, Schuhmacher JS, Altegoer F, Fage CD, Srinivasan V, Linne U, et al. Catalytic mechanism and allosteric regulation of an oligomeric (p)ppGpp synthetase by an alarmone. Proc Natl Acad Sci U S A. 2015;112(43):13348–53.

15. Hogg T, Mechold U, Malke H, Cashel M, Hilgenfeld R. Conformational antagonism between opposing active sites in a bifunctional RelA/SpoT homolog modulates (p)ppGpp metabolism during the stringent response. Cell. 2004;117:57–68.

16. Manav MC, Beljantseva J, Bojer MS, Tenson T, Ingmer H, Hauryliuk V, et al. Structural basis for (p)ppGpp synthesis by the Staphylococcus aureus small alarmone synthetase RelP. J Biol Chem. 2018;293(9):3254–64.

17. Srivatsan A, Han Y, Peng J, Tehranchi AK, Gibbs R, Wang JD, et al. High-precision, whole-genome sequencing of laboratory strains facilitates genetic studies. PLoS Genet. 2008;4(8):e1000139.

18. Yang J, Anderson BW, Turdiev A, Turdiev H, Stevenson DM, Amador-Noguez D, et al. The nucleotide pGpp acts as a third alarmone in Bacillus, with functions distinct from those of (p) ppGpp. Nat Commun. 2020;11(1):5388.

19. Petchiappan A, Naik SY, Chatterji D. RelZ-Mediated Stress Response in Mycobacterium smegmatis: pGpp Synthesis and Its Regulation. J Bacteriol. 2020;202(2).

20. Libby EA, Reuveni S, Dworkin J. Multisite phosphorylation drives phenotypic variation in (p)ppGpp synthetase-dependent antibiotic tolerance. Nat Commun. 2019;10(1):5133.

21. Cao M, Wang T, Ye R, Helmann JD. Antibiotics that inhibit cell wall biosynthesis induce expression of the Bacillus subtilis sigma(W) and sigma(M) regulons. Mol Microbiol. 2002;45(5):1267–76.

22. Czarny TL, Perri AL, French S, Brown ED. Discovery of novel cell wall-active compounds using P ywaC, a sensitive reporter of cell wall stress, in the model gram-positive bacterium Bacillus subtilis. Antimicrob Agents Chemother. 2014;58(6):3261–9.

23. Ackermann M. A functional perspective on phenotypic heterogeneity in microorganisms. Nat Rev Microbiol. 2015;13(8):497–508.

24. Parker DJ, Lalanne JB, Kimura S, Johnson GE, Waldor MK, Li GW. Growth-Optimized Aminoacyl-tRNA Synthetase Levels Prevent Maximal tRNA Charging. Cell Syst. 2020;11(2):121– 30 e6.

25. Dubrac S, Bisicchia P, Devine KM, Msadek T. A matter of life and death: cell wall homeostasis and the WalKR (YycGF) essential signal transduction pathway. Mol Microbiol. 2008;70(6):1307– 22.

26. Kaur P, Rausch M, Malakar B, Watson U, Damle NP, Chawla Y, et al. LipidII interaction with specific residues of Mycobacterium tuberculosis PknB extracytoplasmic domain governs its optimal activation. Nat Commun. 2019;10(1):1231.

27. Beljantseva J, Kudrin P, Andresen L, Shingler V, Atkinson GC, Tenson T, et al. Negative allosteric regulation of Enterococcus faecalis small alarmone synthetase RelQ by single-stranded RNA. Proc Natl Acad Sci U S A. 2017;114(14):3726–31.

28. Takada H, Roghanian M, Caballero-Montes J, Van Nerom K, Jimmy S, Kudrin P, et al. Ribosome association primes the stringent factor Rel for tRNA-dependent locking in the A-site and activation of (p)ppGpp synthesis. Nucleic Acids Res. 2021;49(1):444–57.

29. Gaca AO, Kudrin P, Colomer-Winter C, Beljantseva J, Liu K, Anderson B, et al. From (p)ppGpp to (pp)pGpp: Characterization of Regulatory Effects of pGpp Synthesized by the Small Alarmone Synthetase of Enterococcus faecalis. J Bacteriol. 2015;197(18):2908–19.

30. Yang N, Xie S, Tang NY, Choi MY, Wang Y, Watt RM. The Ps and Qs of alarmone synthesis in Staphylococcus aureus. PLoS One. 2019;14(10):e0213630.

31. Horvatek P, Salzer A, Hanna AMF, Gratani FL, Keinhorster D, Korn N, et al. Inducible expression of (pp)pGpp synthetases in Staphylococcus aureus is associated with activation of stress response genes. PLoS Genet. 2020;16(12):e1009282.

32. Mechold U, Potrykus K, Murphy H, Murakami KS, Cashel M. Differential regulation by ppGpp versus pppGpp in Escherichia coli. Nucleic Acids Res. 2013;41(12):6175–89.

33. Anderson BW, Hao A, Satyshur KA, Keck JL, Wang JD. Molecular Mechanism of Regulation of the Purine Salvage Enzyme XPRT by the Alarmones pppGpp, ppGpp, and pGpp. J Mol Biol. 2020;432(14):4108–26.

34. Vinogradova DS, Zegarra V, Maksimova E, Nakamoto JA, Kasatsky P, Paleskava A, et al. How the initiating ribosome copes with ppGpp to translate mRNAs. PLoS Biol. 2020;18(1):e3000593.

35. Rittershaus ES, Baek SH, Sassetti CM. The normalcy of dormancy: common themes in microbial quiescence. Cell Host Microbe. 2013;13(6):643–51.

36. Bergkessel M, Basta DW, Newman DK. The physiology of growth arrest: uniting molecular and environmental microbiology. Nat Rev Microbiol. 2016;14(9):549–62.

37. Rosenthal AZ, Qi Y, Hormoz S, Park J, Li SH, Elowitz MB. Metabolic interactions between dynamic bacterial subpopulations. Elife. 2018;7.

38. Veening JW, Stewart EJ, Berngruber TW, Taddei F, Kuipers OP, Hamoen LW. Bet-hedging and epigenetic inheritance in bacterial cell development. Proc Natl Acad Sci U S A. 2008;105(11):4393–8.

39. Maamar H, Raj A, Dubnau D. Noise in gene expression determines cell fate in Bacillus subtilis. Science. 2007;317(5837):526–9.

40. Ababneh QO, Herman JK. RelA inhibits Bacillus subtilis motility and chaining. J Bacteriol. 2015;197(1):128–37.

41. Mirouze N, Desai Y, Raj A, Dubnau D. Spo0A∼P imposes a temporal gate for the bimodal expression of competence in Bacillus subtilis. PLoS Genet. 2012;8(3):e1002586.

